# Species-specific evolution of conserved noncoding elements associated with distinctive traits of *Caenorhabditis inopinata*

**DOI:** 10.1101/2024.09.18.613604

**Authors:** Katsunori Tamagawa, Shun Oomura, Asako Sugimoto, Takashi Makino

**Affiliations:** Laboratory of Evolutionary Genomics, Graduate School of Life Sciences, Tohoku University, Aoba-ku, Sendai, Japan; Department of Marine Bioscience, Atmosphere and Ocean Research Institute, The University of Tokyo, 5-1-5 Kashiwanoha, Kashiwa, Chiba, Japan; Department of Integrated Biosciences, Graduate School of Frontier Sciences, The University of Tokyo, 5-1-5 Kashiwanoha, Kashiwa, Chiba, Japan; Laboratory of Developmental Dynamics, Graduate School of Life Sciences, Tohoku University, Aoba-ku, Sendai, Japan

**Keywords:** nematode, conserved noncoding elements, accelerated evolution, body size, *Caenorhabditis inopinata*

## Abstract

Phenotypic evolution is driven by genetic mutations occurring in both protein-coding and noncoding regions of the genome. Conserved noncoding elements (CNEs) have—at least partially—gene regulatory functions and contribute to the evolution of organisms by altering their gene expression. Evolutionary changes in CNEs, including their loss and accelerated evolution, can play crucial roles in shaping species-specific traits. The extensive functional genomic information available for the model nematode *Caenorhabditis elegans*, together with the recent accumulation of high-quality genome sequences from related species, has made *Caenorhabditis* nematodes a powerful system for comparative genomics focused on CNE evolution. The recently described species *Caenorhabditis inopinata* has several peculiar traits notable for its unusually large body size and is regarded as an appropriate species that links phenotypic evolution with genomic changes. Here, using comparative genomics and transcriptomics in *C. inopinata*, we analyzed the evolution of CNEs and inferred their influence during phenotypic evolution. We detected substantial evolutionary changes in CNEs in *C. inopinata* compared to other relatives—changes frequently associated with body morphology and behavior corresponding to distinct ecological traits. Our findings suggest that loss and accelerated evolution of CNE are associated with species-specific traits and provide new insight into the impact of noncoding elements on evolution.

**Significance statement:** Large-scale phenotypic evolution has been observed across a wide range of taxa and may be accompanied by substantial genomic changes. *Caenorhabditis inopinata*, a nematode species closely related to *C. elegans*, shows several distinctive morphological and behavioral traits compared with its congeners, and through comparative genomics and transcriptomics, we identify substantial evolutionary changes in conserved noncoding elements (CNEs)— including their loss and accelerated sequence evolution. Furthermore, these changes are frequently associated with genes involved in morphology and behavior, suggesting a potential link between CNE evolution and species-specific phenotypes.

## Introduction

Previously, the nonprotein-coding elements in genomic sequences were often considered nonfunctional because they did not encode proteins (Alexander et al. 2010). At present, those elements are frequently considered key players in extensive biological phenomena (Siepel et al. 2005; Polychronopoulos et al. 2017). Some noncoding elements are evolutionarily conserved across species—referred to as conserved noncoding elements (CNEs)—and have been shown to play essential roles in gene expression regulation (Siepel et al. 2005; Polychronopoulos et al. 2017). Mutations in those elements cause phenotypic changes, including human diseases (Polychronopoulos et al. 2017). Consequently, in evolutionary biology, the function of noncoding elements in lineage-specific phenotypes is a current focus and is an active area of research. For example, loss of CNEs is frequently observed in the seahorse genome compared to other teleosts (Lin et al. 2016), and accelerated evolution of CNEs is associated with loss of flight in birds (Sackton et al. 2019). In addition to vertebrates, the evolutionary changes in CNEs are associated with the gain and loss of traits of invertebrate lineages (Wang et al. 2015; Rubin et al. 2019; Gonzalez et al. 2024).

Recently, the genome sequence of *Caenorhabditis* nematodes—congeneric with the well-studied model organism *C. elegans*—has rapidly accumulated due to advances in sequencing technologies. The availability of detailed genomic information for *C. elegans*, combined with its compact genome size, make them suitable for comparative genomics (Corsi et al. 2015). In addition to the growing number of sequenced genomes, fascinating new species have been reported, the most striking example being *Caenorhabditis inopinata,* a sibling species of *C. elegans,* isolated from fig syconia in Okinawa, Japan (Kanzaki et al. 2018). Although nematodes are typically characterized by highly constrained morphological evolution, *C. inopinata* markedly differs from its close relatives in body size, behavior, and habitat (Kanzaki et al. 2018; Woodruff et al. 2018). For instance, *C. inopinata* has a long and slender body more than twice the length of other relative species (Kanzaki et al. 2018). Moreover, *C. inopinata* was isolated from fresh fig syconia of *Ficus septica,* whereas other *Caenorhabditis* relatives generally inhabit soil, rotting organic materials, and leaf-litter environments (Kiontke et al. 2011; Kanzaki et al. 2018). Reports of unique genomic features—such as desilencing and accumulation of transposable elements—suggest that these genomic changes may underpin the unusual ecological characteristics of *C. inopinata* (Kanzaki et al. 2018; Hatanaka et al. 2024). Various genetic tools have now been adopted from sibling species of *C. elegans* (Oomura et al. 2022). As a result, *C. inopinata* has emerged as a promising model system for studying phenotypic evolution.

The body size of *C. inopinata* is twice as large as that of *C. elegans* despite their close relationship as sibling species (Kanzaki et al. 2018; Woodruff et al. 2018) (Fig. 1A). In these species, body size reflects growth rate during the transition from larval to adult stages—particularly at the L4 stage—and depends on cell size rather than cell number (Woodruff et al. 2018). Although genetic and environmental factors influence the body size of animals (Silventoinen 2003; Tuck 2014; Texada et al. 2020), maintenance in the laboratory does not reduce the body size of *C. inopinata*, suggesting that its enlargement is driven by the evolutionary changes in gene regulation rather than the environmental factors in their habitats. In *C. elegans*, there is substantial evidence for genetic pathways regulating body size, including TGFβ and BMP pathways (Tuck 2014). Regulation of cuticle collagen, which envelopes the entire body, is a direct factor constraining postembryonic growth and morphology of nematodes (Page and Johnstone 2007; Tuck 2014). However, the relationship between these genetic factors and body size enlargement in *C. inopinata* remains poorly understood.

**Figure 1.**
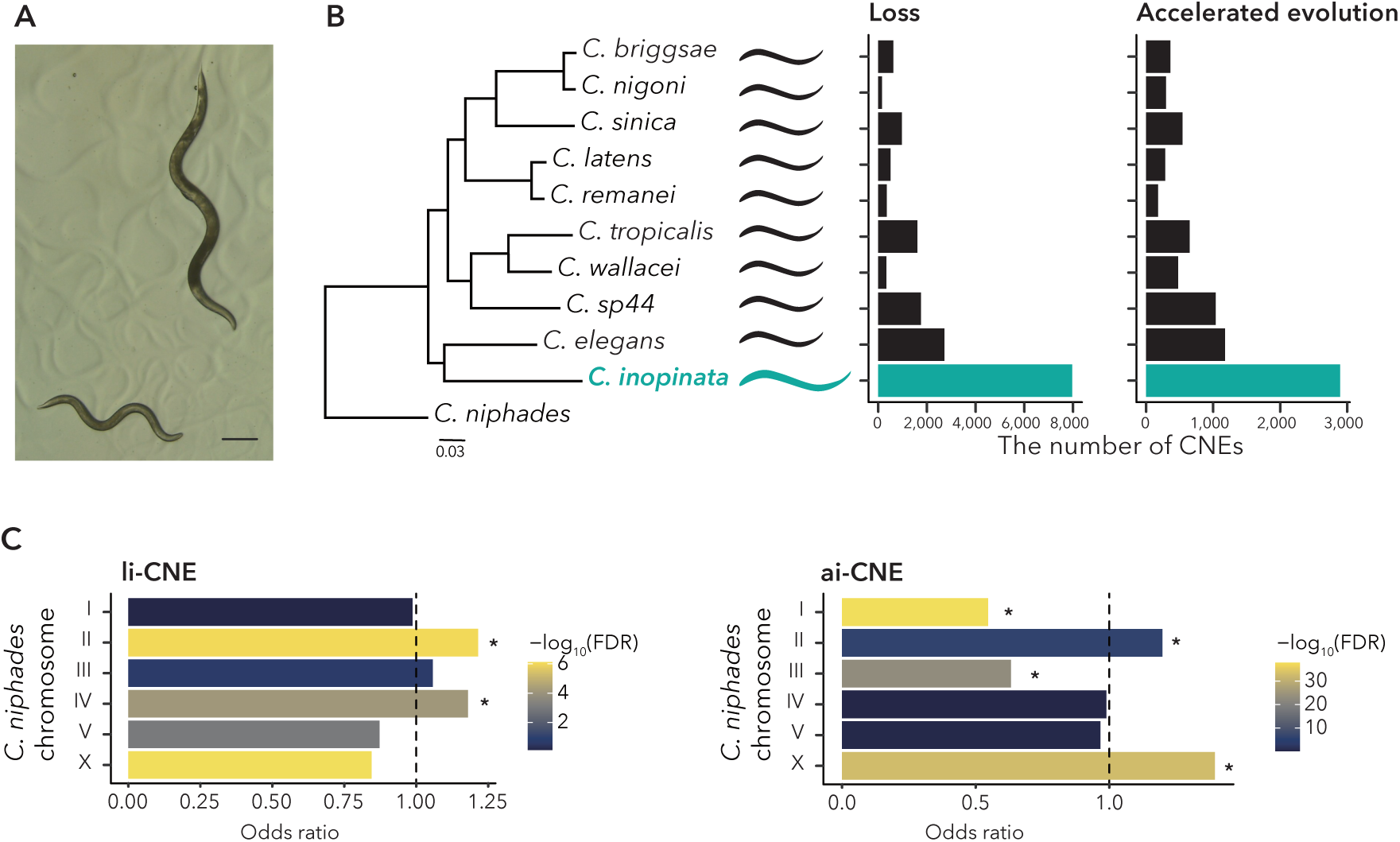
Number and distribution of loss or accelerated evolution of conserved noncoding elements (CNEs). (A) Image of adult of *C. elegans* (lower left) and *C. inopinata* (upper right); scale bar, 200 µm. (B) Numbers of CNE losses and CNEs showing accelerated evolution in each *Caenorhabditis* species. The left panel shows a phylogenetic relationship among *Caenorhabditis* species used in this analysis (adapted from Tamagawa et al. 2024). The middle and right panels indicate the number of CNE losses CNEs with accelerated evolution, respectively. Bars corresponding to *C. inopinata* are highlighted in blue. (C) Chromosomal distribution of CNE loss and accelerated evolution, using *C. niphades* as the reference genome. Asterisks indicate cases in which the observed number of CNE loss or accelerated evolution was greater than expected. Bar colors correspond to the false discovery rate (FDR).

Here, we analyzed CNEs in *C. inopinata* to investigate the evolutionary relationship between noncoding elements and phenotypic traits. Previous analysis showed that the evolution of sexual mode in *Caenorhabditis* nematodes is accompanied by the accelerated evolution of CNEs (Tamagawa et al. 2024). In *C. inopinata,* several CNEs were lost relative to other related species, and accelerated evolution of CNEs was frequently detected. Furthermore, CNEs exhibiting accelerated evolution were significantly enriched near genes associated with body size. These findings support the hypothesis that evolutionary changes in regulatory elements contribute to phenotypic modifications—such as body enlargement—and provide testable candidate genetic factors for tracing the evolutionary trajectory.

## Results

### Number and distribution of li-CNEs and ai-CNEs

Evolutionary changes in CNEs are potentially associated with substantial phenotypic divergence, with CNE loss representing one possible mechanism. As expected, we found twice as many species-specific losses of CNEs in *C. inopinata*, *i.e.,* li-CNEs, as we did in other *Caenorhabditis* nematodes (Grubbs’ outlier test, *p* < 0.01; Fig. 1B middle, Table S1). Accelerated evolution of CNEs is another potential source of phenotypic divergence. We therefore examined CNEs exhibiting accelerated evolution in *C. inopinata*, *i.e.,* ai-CNE, and found a significantly greater number of species-specific ai-CNEs than in other species (Grubbs’ outlier test, p < 0.01; Fig. 1B right, Table S1). CNEs were generally concentrated toward the central region of each chromosome, and no strong positional bias along chromosomes was observed for li- and ai-CNE distribution (Fig. S1, Table S2, 3). However, we detected significant chromosome-specific differences in the distribution of li- and ai-CNEs (Fig. 1C). Whereas li-CNEs were abundantly detected in chromosomes I and IV, ai-CNEs were enriched in chromosomes II and V and were depleted in chromosomes I and III (FDR < 0.05). The neighboring genes of li- and ai-CNE were 3,133 and 1,894, respectively (Table S4, 5). Although these evolutionary changes did not always co-occur in the same neighboring genes, we identified several genes that were neighboring to li-and ai-CNEs when we used only one-to-one orthologs between *C. elegans* and *C. inopinata* to eliminate the effect of gene duplication and loss (Fig. S2).

### Overlapping regulatory elements and neighboring gene functions

Because CNEs are frequently associated with regulatory elements, including promoters and enhancers, we examined the association between li- and ai-CNEs and the regulatory regions based on the genomic annotation of *C. elegans*. li-CNEs significantly overlapped with domains corresponding to enhancers and transcription factor binding sites (TFBS) of the *C. elegans* genome but not with promoter regions (Fisher’s exact test, odds ratio [OR] = 1.26, *p* < 0.01 for enhancer; OR = 1.24, *p* < 0.01 for TFBS; OR = 0.80, *p* = 0.9939 for promoter). We found no significant overlap for 5′ UTR, and li-CNE significantly depleted in 3′ UTR (Fisher’s exact test, OR = 0.92, *p* = 0.1245 for 5′ UTR; OR = 0.5998498, *p* < 0.01 for 3′ UTR). For each regulatory element, ai-CNEs exhibited a pattern similar to that observed for li-CNEs (Fisher’s exact test, OR = 1.23, *p* = 0.0228 for enhancer; OR = 1.85, *p* < 0.01 for TFBS; OR = 1.22, *p* = 0. 1153 for promoter; OR = 0.47, *p* < 0.01 for 3′ UTR; OR = 0.89, *p* = 0.2303 for 5′ UTR).

The genes neighboring li-CNEs were significantly enriched in neuronal expression patterns and several behavioral phenotypes (Fig. 2A, Table S6). Using gene ontology enrichment, we showed the functions associated with neuron and signal transduction (Fig. 2A, Table S6). Similarly, the genes neighboring the ai-CNEs were significantly enriched in neuronal and behavioral phenotypes; in addition, these genes were significantly enriched in the phenotype associated with “body morphology variants” and “small” (Fig. 2B, Table S7). The genes neighboring the ai-CNEs were additionally enriched in “cell part morphogenesis” concerning gene ontology (Fig. 2B, Table S7).

**Figure 2.**
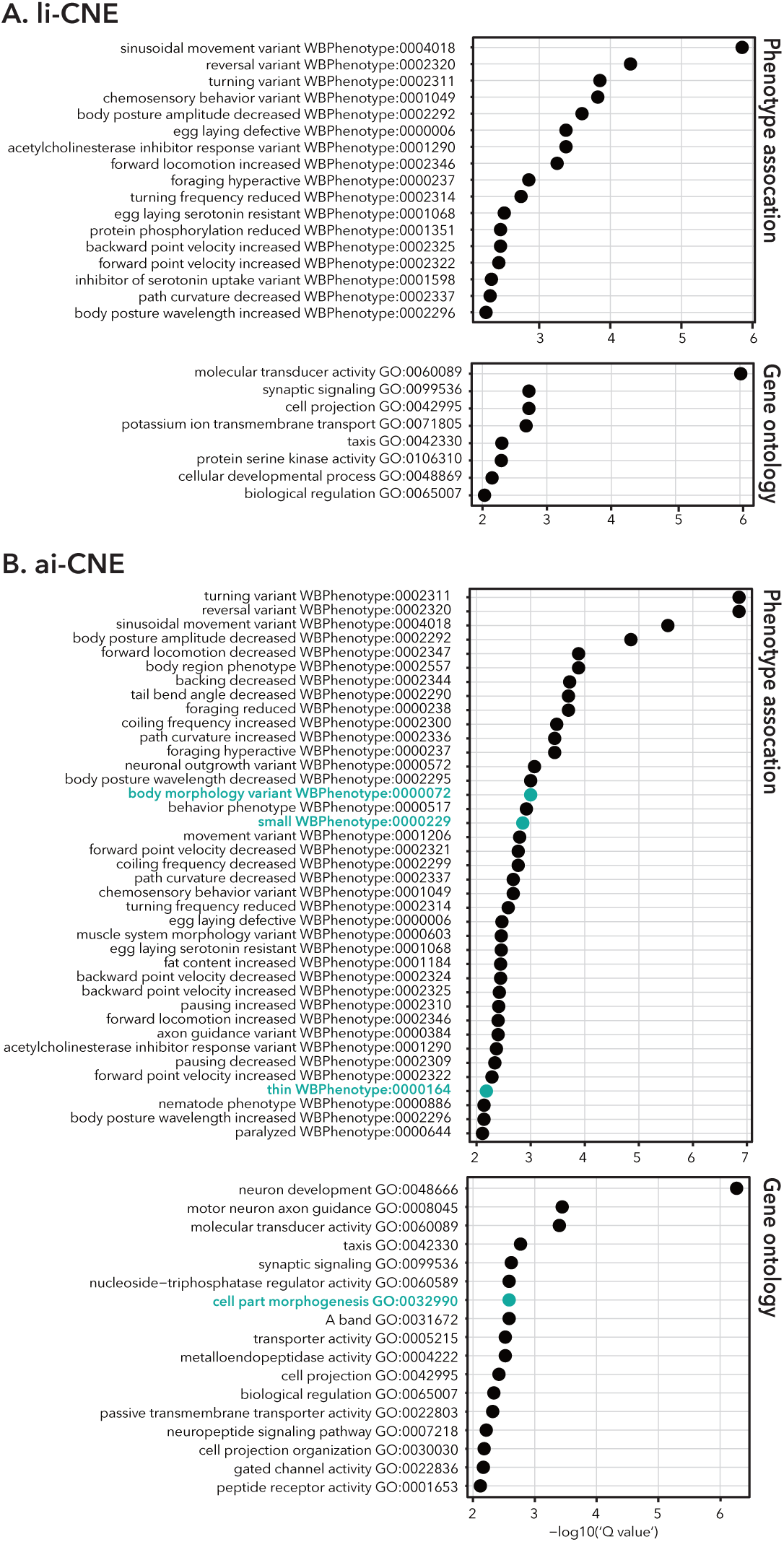
Enrichment analysis of phenotypic and gene ontology terms for genes neighboring conserved noncoding elements (CNEs) (A) Genes neighboring lineage-specific loss of CNEs in *C. inopinata*. (B) Genes neighboring CNEs showing accelerated evolution. Only terms with q-values < 0.05 are shown. Terms associated with body morphology are highlighted in blue.

### Association between gene expression and evolutionary change in CNEs

To clarify the effects of li- and ai-CNEs on the gene expression in *C. inopinata*, we analyzed transcriptomic datasets by comparing gene expression levels between developmental stages within each sex. This analysis identified 7,860 and 8,133 differentially expressed genes (DEGs) in males and females, respectively (Table S8). Genes neighboring li- and ai-CNEs were significantly enriched among genes highly expressed at the L4 stage compared to the YA stage in both males and females (Fig. 3A). By contrast, comparison of gene expression between sexes identified10,106 and 11,356 DEGs at the L4 and YA stages, respectively. Genes neighboring li-and ai-CNEs showed weak or no association with sex-dependent expression at either developmental stage, in stark contrast to their strong association with stage-dependent expression (Fig. 3B). These results suggest that evolutionary changes in CNEs are primarily associated with stage-dependent gene expression. We therefore tested whether this stage-related association was conserved in orthologous genes in sibling species, *C. elegans*. In *C. elegans*, we identified 9,588 and 10,342 DEGs between the L4 and YA stages in males (XO) and feminized hermaphrodite (XX), respectively (Table S9). One-to-one orthologs between *C. inopinata* and *C. elegans* were extracted and classified them into expression trends categories based on their extent relative expression at the L4 or YA stage for each sex (Fig. 3C, Table S10). Genes highly expressed at the L4 stage in both species were frequently adjacent to li- and ai-CNEs, regardless of sex, whereas such enrichment was not observed for genes preferentially expressed at the YA stage. Among genes neighboring li-CNEs, genes highly expressed at the L4 stage in *C. elegans* tended to be highly expressed at the YA stage or to show no differential expression in *C. inopinata* (Fig. 3C left). In contrast, genes neighboring ai-CNEs consistently showed elevated expressed at the L4 stage in *C. inopinata*, *C. elegans*, or both species (Fig. 3C right).

**Figure 3.**
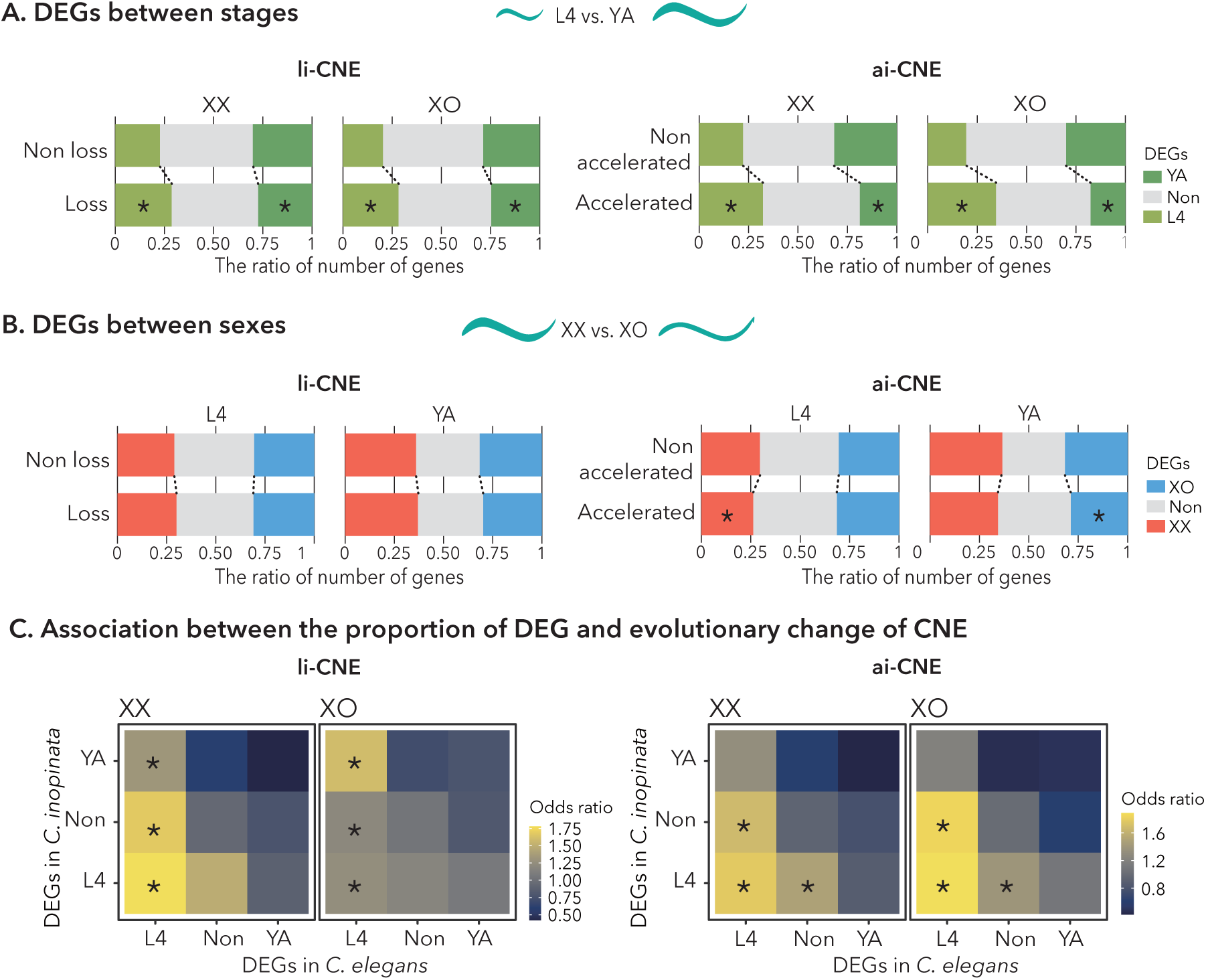
Association between conserved noncoding elements (CNEs) and differential gene expression with respect to developmental stage and sex. (A) Enrichment analysis of genes neighboring each CNEs among differentially expressed genes (DEGs) between the L4 stage and young adult. (B) Enrichment analysis of genes neighboring CNEs among DEGs between females (XX) and males (XO). Upper rows represent genes without loss or accelerated evolution of CNEs, whereas lower rows represent only genes neighboring evolutionarily altered CNEs.

Colors indicate expression bias, and genes without sex-biased expression are labeled “Non.” Asterisks indicate significant differences in DEG proportions between genes with and without li-or ai-CNEs, as determined by a Fisher’s exact test (FDR < 0.05). (C) Cross-species analysis of the association between li- or ai-CNEs and DEGs. Genes were classified according to their expression bias in *C. inopinata* and *C. elegans*. Colors represent the odds ratio for a given gene class being adjacent to li- or ai-CNEs. Asterisks indicate statistical significance (Fisher’s exact test, FDR < 0.05). For example, the lower left rectangle in each panel represents genes that are highly expressed at the L4 stage in both species and are more frequently adjacent to li- or ai-CNEs than expected.

### Effect of evolutionary change in CNEs on interspecies gene expression divergence

To investigate the contribution of li- and ai-CNEs to interspecies changes in gene expression, we compared gene expression levels between the sibling species *C. elegans* and *C. inopinata* (Table S11). In comparisons between females of *C. inopinata* and feminized hermaphrodites of *C. elegans,* genes neighboring li- and ai-CNEs showed higher expression levels in *C. inopinata* than expected based on genome-wide expression distributions (Fig. 4A upper panels). In contrast, in males, genes neighboring li- and ai-CNEs tended to show lower expression levels in *C. inopinata* than expected (Fig. 4A lower panels). Despite these sex-dependent differences in expression patterns, DEGs neighboring ai-CNEs were frequently enriched for functions associated with body size (Fig. 4B, Table S12, 13). In addition, cuticle- and collagen-associated functions were enriched in several comparisons (Fig. 4B, Table S12, 13).

**Figure 4.**
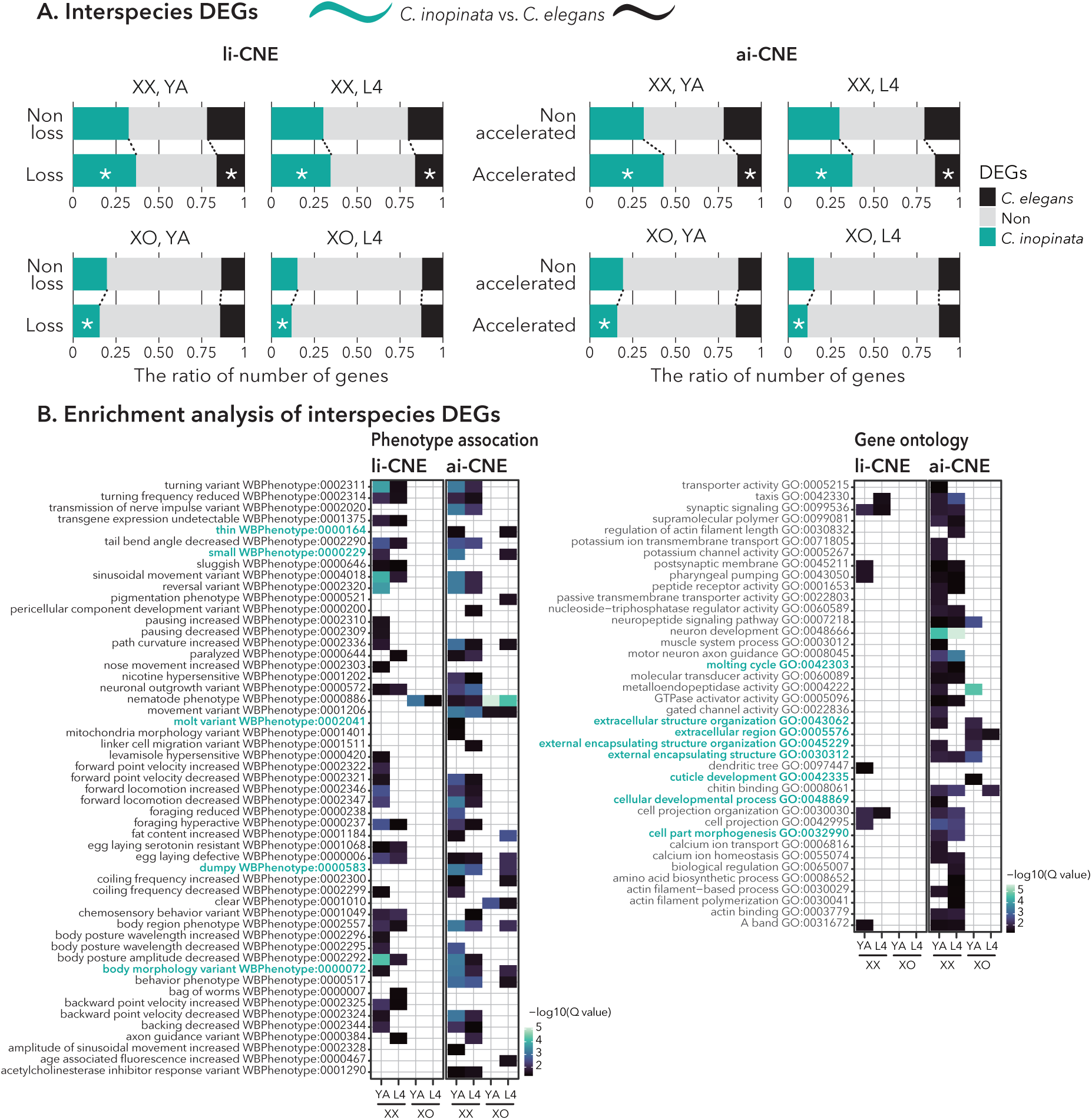
Association between conserved noncoding elements (CNEs) and interspecies differential expression across developmental stage and sex. (A) Enrichment analysis of differentially expressed genes (DEGs) neighboring CNE loss or accelerated evolution between *C. inopinata* and *C. elegans*. The top and bottom panels represent DEG ratio in females of *C. inopinata* compared with feminized *C. elegans* (XX) and in males (XO), respectively. Upper rows show genes without loss or accelerated evolution of CNEs, and lower rows show only genes neighboring CNEs under evolutionary changes. Colors represent the DEG expression bias, and genes with no sex bias are labeled “Non.” Asterisks indicate statistical significance based on Fisher’s exact test (FDR < 0.05). (B) Enrichment analysis of phenotypic associations and gene ontology terms using differentially expressed genes neighboring lost CNEs and CNEs showing accelerated evolution. DEGs were identified for each developmental stage and sex between species. Colors reflect q-values, and only terms with q-value < 0.05 are shown. Terms associated with morphology are highlighted in blue.

### Evolutionary changes in CNEs associated with body size-regulating genes

We identified several li- and ai-CNEs adjacent to genes associated with body size regulation, including collagen and their regulators (Fig. 5). The gene *blmp-1*, which encodes a zinc finger and SET domain-containing proteins, harbored both li- and ai-CNEs in its upstream region (Fig. 5A). *blmp-1* gene expression was high during the L4 stage in most samples, except in XX individuals of *C. elegans* (Fig. 5A right). The cuticular collagen gene *dpy-2* contained ai-CNEs in its proximal upstream region (Figure 5B, C). In *C. inopinata, dpy-2* was expressed at higher levels than in *C. elegans*, particularly at the L4 stage when body size difference between species becomes apparent (Fig. 5B right). In this case, nucleotide substitutions were distributed throughout the entire CNE sequence in *C. inopinata* compared with other species (Fig. 5C).

**Figure 5.**
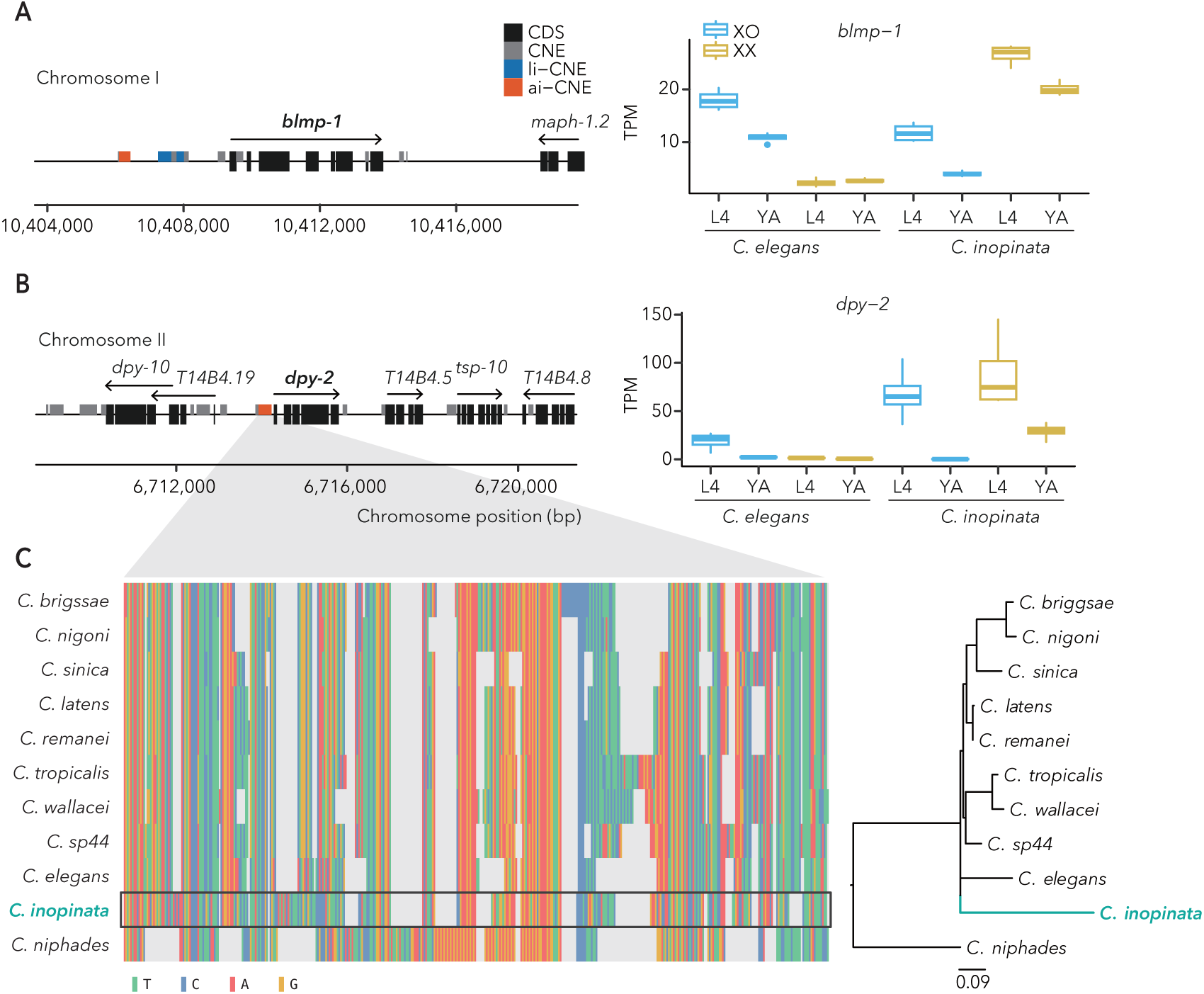
Collagen-related genes neighboring conserved noncoding elements (li- and ai-CNEs) showing loss or accelerated evolution. (A, B) Positional relationships between CNEs and gene expression pattern of *blmp-1* (A) and *dpy-2* (B). Schematics show 5 kbp upstream and downstream of each gene. Arrows over the coding sequences (CDS) indicate the direction of transcription. (C) Nucleotide alignment and phylogenetic tree of ai-CNEs neighboring *dpy-2*. Colors correspond to individual nucleotides.

## Discussion

Genomic changes in noncoding regions have been increasingly recognized as important drivers of phenotypic evolution through their effects on gene regulatory functions (Wang et al. 2015; Lin et al. 2016; Rubin et al. 2019; Sackton et al. 2019). In this study, we demonstrate that evolutionary changes in CNEs including lineage-specific loss or accelerated sequence evolution, occur frequently in the genome of *C. inopinata* and are likely associated with species-specific traits. Both li- and ai-CNEs were significantly associated with changes in gene expression and with evolutionary divergence in the neighboring genes, suggesting a functional link between CNE evolution and regulatory variation. Notably, these associations were particularly evident for genes related to body size regulation, a defining phenotypic feature of *C. inopinata*. Together, our results support the idea that mutations in CNEs contribute substantially to phenotype evolution and provide new insight the regulatory basis underlying the emergence of extraordinary traits of *C. inopinata*.

The loss and accelerated evolution of CNEs in *C. inopinata* were approximately twofold more frequent than in other species (Fig. 1), possibly reflecting the dynamic nature of genome evolution in this lineage. Previous studies have reported a greater abundance of transposable elements (TE) in the *C. inopinata* genome than in other species, and these elements are thought to be derepressed (Kanzaki et al. 2018). Consistent with this view, recent analysis tracking transposon movement under laboratory conditions suggest that TEs remain highly active in *C. inopinata* (Hatanaka et al. 2024). Because TEs are a major source of genomic variation (Chuong et al. 2017; Klein and O’Neill 2018), the extensive changes observed in CNEs may, at least in part, reflect TE activity. The observed depletion of CNE in chromosome arms may also be related to the enrichment of repetitive elements in these regions in *Caenorhabditis* nematodes (Fig.S1) (Woodruff and Teterina 2020). Notably, li- and ai-CNEs tended to be enriched on different chromosomes (Fig. 1C), and the sets of genes neighboring li- and ai-CNEs only partly overlapped (Fig. S2). These observations suggests that CNE loss and accelerated CNE evolution are driven by distinct evolutionary processes. CNE loss may be associated with structural changes such as deletion and insertion, whereas accelerated evolution may be driven by the accumulation of point mutation. Such differences may reflect distinct evolutionary timescale or genomic contexts underlying the emergence of li- and ai-CNE. If li-CNEs originated earlier than ai-CNEs, extensive sequence erosion over evolutionary time could have rendered many ancestral mutations undetectable by current genome comparisons. Differences in the evolutionary timing of CNE loss versus acceleration may therefore be linked to the functional properties, as well as to periods of heightened genomic instability or adaptation to specific environmental conditions.

Evolutionary changes in CNE preferentially overlapped with enhancers and TFBSs, rather than with promoter regions. Because promoters are essential for the initiation of gene expression, evolutionary changes such as loss or accelerated evolution of CNEs may be constrained in promoter regions to preserve their vital function (Andersson 2015). Instead, loss and accelerated evolution of noncoding elements may contributed to the evolution and modulation of spatiotemporal gene regulation through non-promoter regulatory regions, such as enhancers.

Regarding phenotypic associations, genes neighboring li- and ai-CNEs were frequently associated with neuronal development and behavior (Fig. 2). The habitat of *C. inopinata*—inside fresh fig fruits—differs markedly from that of its close relatives, which typically inhabit rotten fruits or soil (Kiontke et al. 2011). This distinct ecological niche may have driven the evolution of adaptive behaviors, such as foraging, dispersal and mating behavior in *C. inopinata*. Notably, previous study have reported a substantial contraction of seven-transmembrane G protein-coupled receptor (7TM-GPCR) gene families in the *C. inopinata* genome, potentially reflecting adaptation to its restricted habitat and close association with fig and their vector insects (Kanzaki et al. 2018). Furthermore, ai-CNEs frequently adjacent to genes regulating body morphology phenotypes (Fig. 2B), suggesting that evolutionary changes in CNEs contribute to the emergence of habitat-specific traits of *C. inopinata* (Kanzaki et al. 2018; Woodruff et al. 2018). Associations with body morphology-related phenotypes, such as “small” and “thin,” were detected exclusively for ai-CNE. This pattern may indicate an accelerated CNE evolution occurred concomitantly with the evolution of body morphology in *C. inopinata*. Such tendency could arise from the acquisition of novel regulatory functions by CNEs through accelerated sequence evolution.

Genes neighboring li- and ai-CNEs were significantly associated with stage-biased expression in *C. inopinata* (Fig. 3A), indicating a close relationship between evolutionary changes in CNEs and developmental regulation of gene expression. Consistent with this pattern, orthologous genes between *C. inopinata* and *C. elegans* showed preferential expression at the larval stage in both species (Fig. 3C), supporting the involvement of li- and ai-CNEs in larval stage gene regulation. Body length of *C. inopinata* depends largely on growth during the transition from larval to adult stages (Woodruff et al. 2018). Therefore, evolutionary changes in CNEs that affect stage-specific gene regulatory programs may contributed to the enlargement of body size in this species. In addition, several genes neighboring li- and ai-CNEs exhibited distinct developmental expression patterns between *C. inopinata* and *C. elegans* (Fig. 3C), suggesting that the evolutionary changes in CNEs may also underlie species-specific shifts in gene regulation. Taken together, these observations imply that loss and accelerated evolution of CNEs are associated with regulatory transitions during development and may have played an important role in phenotypic divergent, including body size enlargement, in *C. inopinata*.

When gene expression was directly compared between *C. inopinata* and *C. elegans*, we observed significant associations between evolutionary change in CNEs and interspecies differential expression (Fig. 4A). Genes neighboring li- and ai-CNEs were frequently associated with functions related to behavior and body morphology, with particularly more pronounced enrichment observed for ai-CNE–associated genes (Fig. 4B). These results suggest that evolutionary change in CNEs may contribute to interspecies shifts in gene expression underlying the emergence of unique traits of *C. inopinata*. Intriguingly, the association between CNE evolution and gene expression displayed clear sex-dependent pattern (Fig. 4). Such sex dependency may reflect difference in how evolutionary changes in CNEs affect regulatory programs in males and females. Because sex determination genes identified in *C. elegans* are partially disrupted in *C. inopinata* (Kanzaki et al. 2018), regulatory network underlying sexual differentiation may also be altered, and evolutionary change in CNEs could be involved in sex differentiation. These sex-dependent patterns may also reflect the differences in the genetic basis for body enlargement between sexes.

Regulation of cuticle collagen is essential for postembryonic growth and body morphology of nematodes (Page and Johnstone 2007; Tuck 2014). In this study, we identified li-and ai-CNEs upstream of *blmp-1* and ai-CNEs upstream of *dpy-2* (Fig. 5), both of which are key regulators of cuticle formation and molting. *blmp-1* regulates the gene expression of collagen and is required for proper cuticle function, making it central to molting and developmental progression (Sandhu et al. 2021; Stec et al. 2021). Null mutation in *blmp-1* disrupt the larval-to-adult transition and lead to misregulation of collagens transcribed during intermolt stages, including *dpy-2* functioning (Wu et al. 2022). Consistently, knockdown of *blmp-1* significantly reduces *dpy-2* expression and results in marked body size reduction (Sandhu et al. 2021; Stec et al. 2021). The ai-CNE located upstream of *dpy-2* shows extensive nucleotide substitutions across its entire length and is conserved among *Caenorhabditis* relatives except for the outgroup species, *C. niphades* (Fig. 5C). This pattern suggests that the ai-CNE was acquired after the divergence from *C. niphades* and subsequently accumulated substitutions specifically in the *C. inopinata* lineage, although contributions from additional evolutionary processes cannot be excluded. Taken together, evolutionary changes in CNEs neighboring *blmp-1* and *dpy-2* are likely to alter developmental regulation of cuticle collagen expression and thereby contribute to postembryonic growth dynamics. These regulatory changes provide a plausible molecular basis for the pronounced body size enlargement observed in *C. inopinata*.

Our analysis highlights a link between the evolutionary changes in noncoding regions and large-scale phenotypic evolution in invertebrates, as demonstrated in *Caenorhabditis* nematodes. Accumulating evidence suggests that mutational changes in noncoding elements can have additive effects on phenotype, with individual contributions that are often small to moderate (Pennisi 2017; Polychronopoulos et al. 2017). As a result, disentangling the phenotypic influence of individual evolutionary change in CNEs is remains challenging. In addition, experimental validation of CNE evolution in *C. inopinata* is currently limited by the lack of well-established genome editing methods. Addressing these limitations will require future studies that combine improved genetic tools with targeted functional assays to directly assess the contribution of evolutionary changes in CNEs to phenotype evolution.

## Conclusion

Noncoding elements conserved among species play important roles in gene expression regulation through diverse mechanisms. The comparative genome analysis revealed substantial loss and accelerated evolution of CNE in *C. inopinata*, a species with a unique habitat and large body size, compared to closely related species . These evolutionary changes in CNEs were frequently observed in the vicinity of genes associated with body morphology and behavior. In addition, transcriptomic analysis revealed that both loss and accelerated evolution of CNE are closely associated with developmentally regulated gene expression, particularly during stages relevant to body size enlargement. Together, our results support the view that evolutionary changes in CNE contribute to the acquisition of species-specific traits by modifying gene regulatory programs.

## Materials and Methods

### Whole genome alignment and CNE detection

In this analysis, we used the CNE datasets described by Tamagawa et al. (2024), comprising 133,541 CNEs identified from whole-genome alignment of 11 *Caenorhabditis* species: *C. briggsae*, *C. elegans*, *C. latens*, *C. nigoni*, *C. remanei*, *C. sinica*, *C. tropicalis* (obtained from WormBase ParaSite version WBPS15), *C. niphades* (Sun et al. 2022), *C. inopinata* (GCA_003052745.1), *C. sp44*, and *C. wallacei* (http://caenorhabditis.org/) using Progressive Cactus (Armstrong et al. 2020). For genes with multiple isoforms, we retained the longest isoforms. We aligned single-copy orthologs (4,930 genes) using mafft (Katoh et al. 2002) and removed poorly aligned regions by trimAl with default parameters (Capella-Gutiérrez et al. 2009). We estimated the consensus species tree from the protein alignment by maximum likelihood with 1,000× bootstrapping using IQ-tree with automatic substitution model selection (Nguyen et al. 2015). With *C. niphades* as a reference, we processed the genome alignment using maftools (Mayakonda et al. 2018) and detected conserved sequences using PhastCons per instructions (Siepel et al. 2005). We classified conserved sequences not overlapping with protein-coding sequences using BEDtools (Quinlan and Hall 2010) and defined sequences longer than 10 bp in the *C. niphades* genome as CNEs.

### Evolutionary change in CNEs and association with neighboring gene function and regulatory elements

We defined the loss of a CNE in *C. inopinata* (li-CNE) as CNEs that are aligned in all other species but not aligned in *C. inopinata*. Next, we detected CNEs exhibiting accelerated evolution in the *C. inopinata* lineage (ai-CNEs) using the log-likelihood ratio test implemented in phyloP with default parameters (Pollard et al. 2010). We removed the CNE sequences that were redundantly aligned or more than twice as long as the corresponding *C. niphades* sequence. To determine whether the number of li- or ai-CNEs in *C. inopinata* was an outlier, we conducted Grubbs’ test using the statistical package of R, “outliers” (cran.r-project.org/web/packages/outliers). We tested for bias in the distribution of the number of li- or ai-CNEs among chromosomes using Fisher’s exact test with R for each chromosome. To perform enrichment analysis of neighboring genes of loss or accelerated evolution of CNEs using WormBase Enrichment Suite (Angeles-Albores et al. 2018), we extracted one-to-one orthologs (11,246 genes) between *C. elegans* and *C. inopinata*. For all gene set enrichment analyses, we used only one-to-one orthologs as background and target gene sets. We also tested the association between the accelerated evolution of CNEs and each regulatory region in the *C. elegans* genome registered in WormBase using Fisher’s exact test with R.

### Gene expression analysis between stage, sex, and species

We downloaded all Illumina paired-end raw reads generated by Tamagawa et al. (2024) for each developmental stage—L4 and young adult (YA)—and sex—male (XO) and feminized hermaphrodite/female (XX)—of *C. elegans* and *C. inopinata* from the SRA (BioProject identifier PRJDB14254). In this RNA-seq dataset, *fem-1* (*hc17*) mutant worms were used as feminized hermaphrodites of *C. elegans*. We quantified the mRNA abundance for each sample using RSEM (Li and Dewey 2011), read mapping performed by Bowtie2 (Langmead and Salzberg 2012) using default parameters. We conducted pairwise comparisons of gene expression levels using the Wald test in DESeq2 v1.36.0 (Love et al. 2014). For this analysis, we filtered out genes with very low expression (sum of expected count < 10). We detected genes with a false discovery rate (FDR) of < 0.01 as differentially expressed genes (DEGs) and determined the association between li- or ai-CNEs and DEGs between stage and sex in *C. inopinata* using a one-sided Fisher’s exact test. To examine the association between li- or ai-CNEs and gene expression changes between species, we classified one-to-one orthologs based on expression patterns between L4 and YA in each species. The resulting expression classifications were evaluated using a one-sided Fisher’s exact test. To assess whether li- or ai-CNEs are linked to interspecies DEGs, we performed a one-sided Fisher’s exact test as described above. We conducted the enrichment analysis of interspecies DEGs neighboring li- or ai-CNEs using WormBase Enrichment Suite (Angeles-Albores et al. 2018).

## Declarations

### Ethics approval and consent to participate

Not applicable.

### Consent for publication

Not applicable.

### Data Availability

All data are available in the Supplementary Materials, and Dryad (https://doi.org/10.5061/dryad.fxpnvx15t).

## Authors’ contributions

K.T., S.O. and T.M. designed the study. K.T. conducted all analyses. K.T., S.O., A.S. and T.M. wrote and reviewed the manuscript.

## Acknowledgements

Computations were partially performed on the NIG supercomputer at ROIS National Institute of Genetics.

