## Supplementary figures for "Species-specific evolution of conserved noncoding elements associated with distinctive traits of *Caenorhabditis inopinata*"

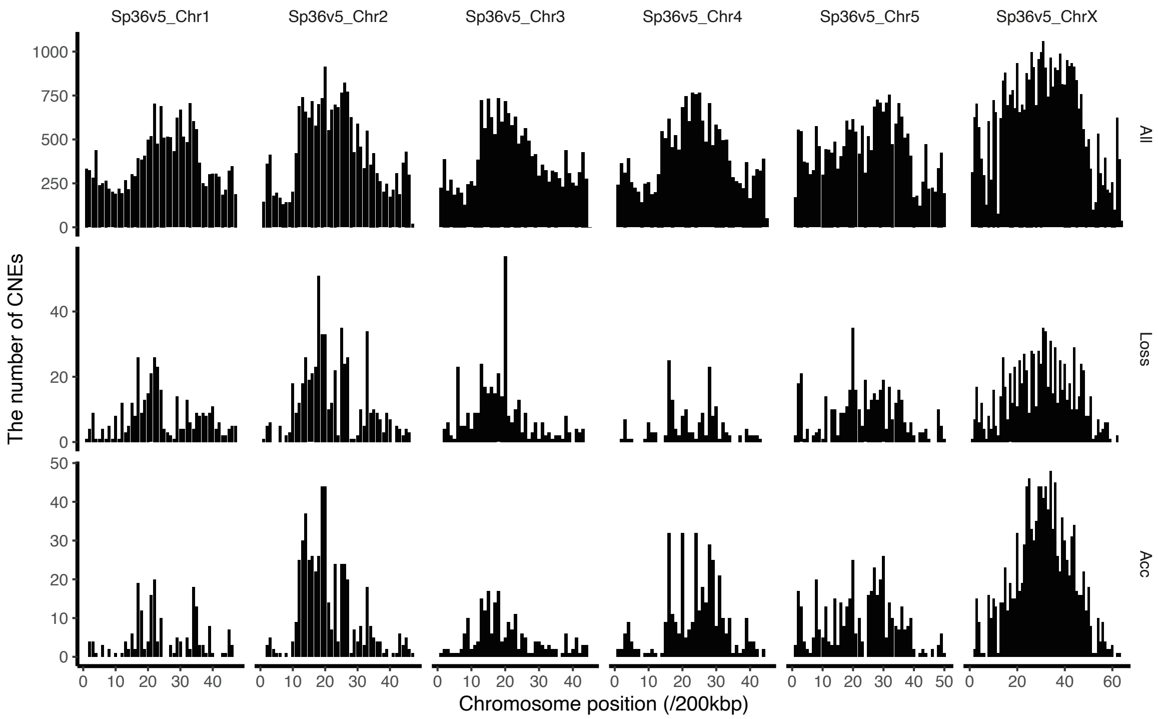


**Figure S1. Chromosomal distribution of conserved noncoding elements (CNEs),** as well as loss and accelerated evolution of CNEs, **in *C. niphades* genome.** The number of CNE is counted within the 200-kbp windows along each chromosome.

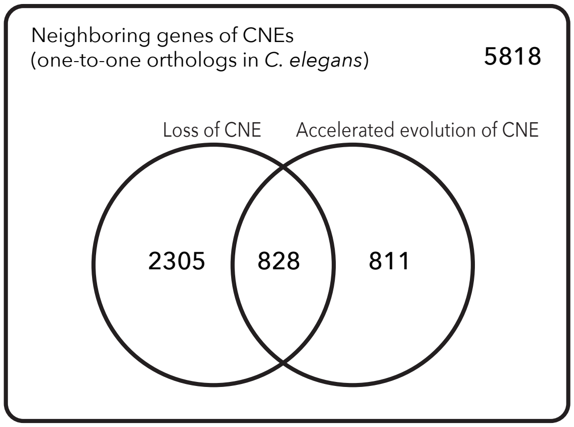


**Figure S2. Overlap of neighboring genes associated with loss and accelerated evolution of conserved noncoding elements (CNEs).**
